# The frontal aslant tract (FAT) and its role in speech, language and executive function

**DOI:** 10.1101/249912

**Authors:** Anthony Steven Dick, Dea Garic, Paulo Graziano, Pascale Tremblay

**Author notes:** Corresponding Author: Anthony Steven Dick, Ph.D., Associate Professor, Director, Cognitive Neuroscience Program and Graduate Certificate in Cognitive Neuroscience, Department of Psychology, Florida International University Modesto A. Maidique Campus AHC4 454, 11200 S.W. 8th Street, Miami, FL 33199; Lab; Fx, Webpage: http://dcn.fiu.edu.

## Abstract

In this review, we examine the structural connectivity of a recently-identified fiber pathway, the frontal aslant tract (FAT), and explore its function. We first review structural connectivity studies using tract-tracing methods in non-human primates, and diffusion-weighted imaging and electrostimulation in humans. These studies suggest a monosynaptic connection exists between the lateral inferior frontal gyrus and the pre-supplementary and supplementary motor areas of the medial superior frontal gyrus. This connection is termed the FAT. We then review research on the left FAT’s putative role in supporting speech and language function, with particular focus on speech initiation, stuttering and verbal fluency. Next, we review research on the right FAT’s putative role supporting executive function, namely inhibitory control and conflict monitoring for action. We summarize the extant body of empirical work by suggesting that the FAT plays a domain general role in the planning, timing, and coordination of sequential motor movements through the resolution of competition among potential motor plans. However, we also propose some domain specialization across the hemispheres. On the left hemisphere, the circuit is proposed to be specialized for speech actions. On the right hemisphere, the circuit is proposed to be specialized for general action control of the organism, especially in the visuo-spatial domain. We close the review with a discussion of the clinical significance of the FAT, and suggestions for further research on the pathway.

**Highlights:**

- The frontal aslant tract (FAT) is a recently identified fiber pathway
- It connects inferior frontal gyrus with medial frontal motor areas
- The left FAT has been associated with speech and language function
- The right FAT has been associated with inhibitory control
- Both FAT pathways may function in sequential motor planning

## The frontal aslant tract (FAT) and its role in speech, language and executive function

The advent of diffusion-weighted magnetic resonance imaging (DW-MRI) has led to an increased interest in accomplishing one of the fundamental goals of human neuroscience—the comprehensive mapping of the cerebral white matter of the brain. It is these short- and long-range axonal connections that comprise the “human connectome,” or “wiring” of the brain, and an understanding of their anatomical connectivity and functional associations is important for establishing a complete model of brain function. Much of this work began in the late 19^th^ and early 20^th^ centuries, with detailed investigations of the white matter of the brain, most notably in the seminal work of Déjèrine (Déjèrine, 1895, 1901) and Flechsig (1920) using histological staining methods. For the most part, DW-MRI has reinforced these definitions of fiber pathways, and additional ones that were delineated with post-mortem methods, such as blunt fiber dissection (Krieg, 1957; Ludwig & Klingler, 1956; Rosett, 1933).

However, DW-MRI has also led to the definition of new fiber pathways. One of these fiber pathways is the frontal aslant tract (FAT), which has only been identified in the last decade. Although noted earlier (Ford, McGregor, Case, Crosson, & White, 2010; Lawes et al., 2008; Oishi et al., 2008), Catani, Theibaut-de Schotten and colleagues (Catani et al., 2012; Thiebaut de Schotten, Dell’Acqua, Valabregue, & Catani, 2012) defined the pathway and coined the term “aslant tract” due to its oblique course in the frontal white matter. It has now become fairly established that such a pathway exists, and that it connects the posterior inferior frontal gyrus (IFG) with medial aspects of the frontal lobe in the superior frontal gyrus and cingulate gyrus and sulcus—namely the pre-supplementary motor area (pre-SMA), supplementary motor area (SMA), and anterior cingulate cortex (Figure 1).

**Figure 1:**
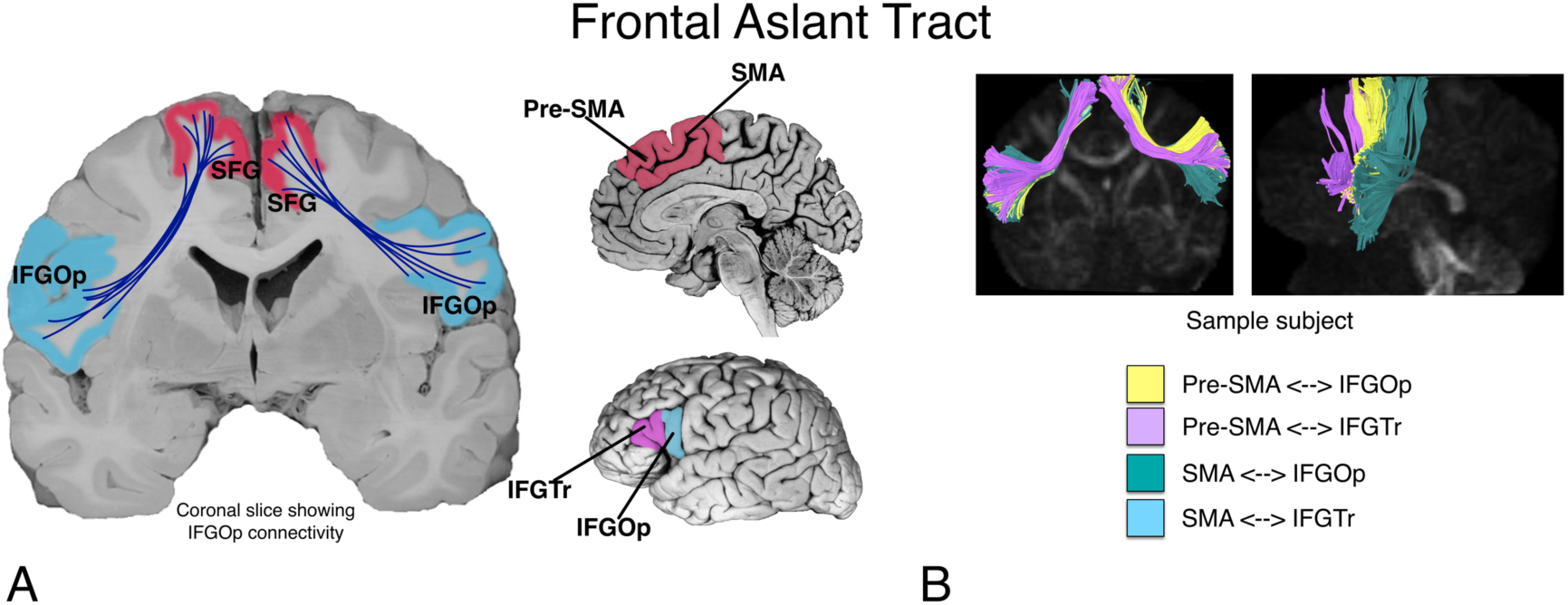
A. The putative connectivity between inferior frontal gyrus (*pars opercularis*; IFGOp, and *pars triangularis*; IFGTr) and pre-supplementary and supplementary motor area (pre-SMA and SMA) in the medial superior frontal gyrus (SFG), supported by the frontal aslant tract (FAT). Figure modified from Dick, A. S., Bernal, B., & Tremblay, P. (2014). The language connectome: new pathways, new concepts. *Neuroscientist, 20*(5), 453-467. B. A sample subject showing four subcomponents of the FAT.

In this review, we strive toward two goals. First, we attempt to establish, based on the available literature, the putative connectivity of the FAT. Second, we attempt to establish the putative functional associations of the FAT, in both the left and the right hemispheres. In the first section, we address the known connectivity of the tract, as well as potential uncertainties. In the second section, we address the functional associations of the left FAT. In the third section, we explore functional associations of the right FAT. In the final section, we propose a model of the function of the FAT in both hemispheres, with some speculation about the clinical significance of the tract and areas of future research.

## 1. Anatomy and connectivity of the Frontal Aslant Tract

Although its description in humans is relatively recent, connections between the lateral inferior frontal cortex and medial superior frontal cortex have been described previously in non-human primates. For example, Thiebaut de Schotten and colleagues (Thiebaut de Schotten et al., 2012) note the similarity of the human FAT with a fiber pathway described in a single macaque studied with autoradiography. This fiber pathway, reported in Case 25 of Schmahmann and Pandya (2006) is similar to, though not identical to, the human FAT. Namely, the injection site is reported to be in the face area of the precentral gyrus, in Brodman Area 4 (i.e., motor cortex). Some terminations from this injection site are reported in the SMA, but not pre-SMA, of the superior frontal gyrus. In some respects, this termination is not surprising given the known connectivity of the SMA to the motor cortex.

More compelling evidence is provided by Petrides and Pandya (2002), who showed that tracer injections in the anterior IFG (namely BA 45 and 47) of six macaques resulted in labeled terminations in the medial superior frontal and cingulate cortex (including pre-SMA). Furthermore, Schmahmann and Pandya (2006) report, in Case 29 of their monograph, that tracer injections into rostral SMA/pre-SMA terminate in area 44 of the IFG. Notably, though, at least one study showed that injections to pre-SMA do not project to area 45 (Wang, Isoda, Matsuzaka, Shima, & Tanji, 2005), and there is no mention of such fibers in the seminal work of Mettler (1935) investigating the fibers of the frontal lobe in the macaque. However, the lack of findings from these latter studies represent a null finding, which should be interpreted with caution, as this may be due to methodological shortcomings. For example, in the case of Mettler, the methods of investigation have been vastly improved since that publication. Where there is a direct attempt to define medial superior frontal and inferior frontal connectivity, for example in the study by Petrides and Pandya (2002), the evidence is present in multiple animals and is therefore more compelling.

The earliest description of this pathway in humans appears in the literature around 2007 and 2008 (we could not find an earlier mention of the pathway in the historical literature from the 19^th^ and 20^th^ centuries). In a 2007 paper, Aron and colleagues (2007) reported connectivity between the pre-SMA and the IFG. Although they did not name it at the time, it is clear that they were identifying the FAT. In another study, Lawes and colleagues (Lawes et al., 2008) conducted an early DW-MRI study combined with post-mortem dissection methods to assess the correspondence between the two methods. In that study, they reported a connection in the DW-MRI analysis between the superior frontal gyrus and the IFG, specifically the *pars triangularis* (IFGTr), and this was verified using post-mortem dissection (albeit on different brains). In the same year, Oishi and colleagues (Oishi et al., 2008) tracked diffusion streamlines from a large superior frontal region of interest (ROI) to an IFG ROI. These streamlines were labeled “frontal short association fibers”. Both tracts defined in these studies contained what we now define to be fibers of the FAT.

In another early DW-MRI study, Ford and colleagues (Ford et al., 2010) used as a point of departure the known inferior frontal-medial superior frontal connectivity described in macaque (Petrides & Pandya, 2002), and explicitly targeted the connections between the IFG and the medial superior frontal cortex in human subjects. They found evidence for connectivity between the posterior IFG and pre-SMA and SMA, and although they did not name the FAT, their description is consistent with the current understanding of the tract. A similar study by Kinoshita and colleagues (Kinoshita et al., 2012), using DW-MRI and blunt fiber dissection in 8 post-mortem brains, also shows what appears to be the FAT. Although this study focused on IFG connections with the lateral superior frontal gyrus, many of the IFG fibers also project to the medial superior frontal gyrus.

Catani, Theibaut-de Schotten and colleagues (Catani et al., 2012; Thiebaut de Schotten et al., 2012) were the first to explicitly name the FAT. They conducted a comprehensive study of the association and U-fiber pathways in the frontal lobe, one of which was the FAT. In this study, they describe the FAT as a pathway that projects predominantly between IFGOp and pre-SMA. The FAT has since been described using blunt fiber dissection techniques in postmortem brains (Goryainov et al., 2017; Koutsarnakis et al., 2017), which adds to the probability that a genuinely new fiber pathway has been described.

Most studies that have followed find that the predominant origin/termination site in the IFG is the *pars opercularis* (Bozkurt et al., 2016), and the predominant connection in the medial superior frontal gyrus is the pre-SMA. However, additional origin/termination paths are also reported. Connectivity with the IFGTr is common, though less consistent than the IFGOp, while reported connections with the more anterior *pars orbitalis* are rare (Szmuda et al., 2017).

Like the inferior frontal connections, connections to and from medial superior frontal cortex are multifaceted. For example, using DW-MRI, Mandelli and colleagues (Mandelli et al., 2014) report connectivity between posterior IFG and pre-SMA/SMA. In another study of eleven post-mortem human brains, using blunt fiber dissection Bozkurt and colleagues (2016) reported that the FAT arises in the anterior SMA and pre-SMA and connects to IFGOp. This was supported by DW-MRI on two participants. Catani and colleagues (Catani et al., 2013) also reported connectivity between the IFGOp and anterior cingulate cortex, along with pre-SMA. In a study of healthy controls and post-mortem subjects, Vergani and colleagues (Vergani et al., 2014) describe SMA connectivity with the *pars opercularis*. Finally, Baker and colleagues (Baker et al., 2018) describe a fiber pathway that they term the “crossed frontal aslant”, which describes connectivity of SMA with homologous and neighboring regions of the contralateral medial and lateral superior frontal cortex, which travel through the anterior corpus callosum. Although they described these fibers as comprising a new pathway, we feel these authors are describing the well-known and already-described contralateral connectivity of the anterior corpus callosum (Schmahmann & Pandya, 2006). We thus believe that it would not be parsimonious to ascribe a new name to these fibers.

One study has reported connectivity between the anterior cingulate gyrus and the anterior insula via the FAT (Y. Li et al., 2016). However, the ROIs in that study were in the medial white matter, not in the cortex. Examination of the terminations of their tracks are instead in the IFG and medial superior frontal gyrus, not in insula or anterior cingulate. That said, a recent mapping of the structural connectivity of subdivisions of the insula suggests that the dorsal anterior insula makes connections with the more anterior superior frontal gyrus (Nomi, Schettini, Broce, Dick, & Uddin, 2017). Using DW-MRI, Mandelli and colleagues (Mandelli et al., 2014) also report connectivity between SMA and anterior insula, which has previously been associated with apraxia of speech (Dronkers, 1996). Electrophysiologic evidence of insular connectivity to pre-SMA and SMA is also available (Enatsu et al., 2016). Some of the fibers supporting this connectivity may travel as part of the FAT.

Finally, a small number of studies have provided electrophysiological evidence of a monosynaptic connection between IFG and medial superior frontal gyrus (Enatsu et al., 2016; Matsumoto et al., 2007; Ookawa et al., 2017; Swann et al., 2012). Enatsu and colleagues (2016) investigated 18 patients using cortico-cortical evoked potentials (CCEP) and found that stimulation of the pre-SMA induces a robust response in IFG. Matsumoto and colleagues (2007) investigated 7 patients with CCEP, with an emphasis on investigating connectivity among areas of the motor system, including pre-SMA and SMA. Some stimulation sites covered the posterior IFG, and indicated connectivity with SMA. These stimulation sites were, however, very close to ventral premotor cortex, which may explain the tendency to connectivity with SMA. Swann and colleagues (2012) conducted a case study using CCEP, electrocorticography (EcOG), and DW-MRI, suggesting direct connectivity between these regions. This study also provided information about the directional nature of information transfer during a stop-signal task and is discussed in more detail in a later section. Briefly, it appears that pre-SMA activity precedes IFG activity during this task, which has implications for models of the FAT’s role in stopping motor responses. Finally, using CCEP and DW-MRI in 8 adult patients, Ookawa and colleagues (2017) showed that stimulation of the IFG elicits a response in mSFG within ∼19-48 ms, on average. Similarly, stimulation of the mSFG elicits a response in IFG within ∼24-70 ms, on average. The latency is significantly shorter for stimulation to the IFG, although both latencies are consistent with a monosynaptic projection between the regions. Thus, the electrophysiologic evidence for IFG and medial superior frontal connectivity is substantial and consistent with the structural findings.

### Summary

There is now ample evidence for a structural connection between inferior frontal and medial superior frontal gyrus (Martino & De Lucas, 2014), which is referred to as the frontal aslant tract (FAT). There is also evidence from electrophysiology to suggest that the tract supports monosynaptic connectivity between these regions. Thus, the FAT connectivity identified initially in DW-MRI has been validated using additional anatomical and electrophysiological methods and is unlikely to represent an artifact of the DW-MRI method.

## 2. Functional Associations of the Left Frontal Aslant Tract in Speech and Language

Given its connectivity with the IFG, which has traditionally been referred to as “Broca’s area”, a region important for speech and language (Tremblay & Dick, 2016), and with the pre-SMA and SMA areas associated with aphasia of the supplementary motor area (Ardila & Lopez, 1984) and with speech production in typical people (Tremblay & Gracco, 2009), it is not surprising that the vast majority of studies on the function of the left FAT have focused on speech and language. Indeed, there is extensive evidence that the left IFG and pre-SMA/SMA are associated with important componential processes in speech and language. Imaging studies have shown that the left IFG is associated with controlled lexical and phonological selection/retrieval in a number of linguistic domains, including in the understanding of sign language and gesture (Badre, Poldrack, Pare-Blagoev, Insler, & Wagner, 2005; Devlin, Matthews, & Rushworth, 2003; A. S. Dick, Mok, Raja Beharelle, Goldin-Meadow, & Small, 2014; Emmorey, Mehta, & Grabowski, 2007; Gough, Nobre, & Devlin, 2005; Katzev, Tuscher, Hennig, Weiller, & Kaller, 2013). The pre-SMA/SMA regions are more associated with motor selection and execution in both speech and non-speech domains (e.g., for manual movements). The pre-SMA is especially thought to play a role in higher-order selection, conflict monitoring and resolution (Tremblay & Gracco, 2006, 2009; Tremblay & Small, 2011), as it does not make a direct connection to the primary motor cortex, the spinal cord, or the cranial nerve motor nuclei (Dum & Strick, 1991; Lu, Preston, & Strick, 1994; Luppino, Matelli, Camarda, Gallese, & Rizzolatti, 1991). Execution of movement may rely more on the SMA and its connections with motor cortex (Tremblay & Gracco, 2009, 2010). These regions are recruited during more complex volitional movements in non-linguistic tasks (Lau, Rogers, & Passingham, 2006; Nachev, Rees, Parton, Kennard, & Husain, 2005; Ullsperger & von Cramon, 2001), including during manual gesture, finger movements, saccades, and notably during tasks involving high response competition/conflict such as task switching (Derrfuss, Brass, & von Cramon, 2004; Mars, Piekema, Coles, Hulstijn, & Toni, 2007; Rushworth, Hadland, Paus, & Sipila, 2002) and flanker tasks (Fan et al., 2007; Nachev et al., 2005; Ullsperger & von Cramon, 2001). In the linguistic domain, volitional word production tasks are associated with a higher activation level than more automatic and more externally constrained tasks (Alario, Chainay, Lehericy, & Cohen, 2006; Etard et al., 2000; Tremblay & Gracco, 2006; Tremblay & Small, 2011).

Damage to these regions is also associated with speech/language disorders. Non-fluent aphasia is a common symptom following lesion to the left perysilvian area, in particular IFG. Despite intense speech therapy, recovery is often incomplete (Kertesz & McCabe, 1977; Pedersen, Jorgensen, Nakayama, Raaschou, & Olsen, 1995; Wade, Hewer, David, & Enderby, 1986). Lesion or tumor resection of the pre-SMA and SMA can also lead to motor and speech deficits, characterized by a global akinesia and hypoflexia, especially for volitional movements and volitional speech. Speech motor deficits are typically more profound on the contralesional side (Bannur & Rajshekhar, 2000; Laplane, Talairach, Meininger, Bancaud, & Orgogozo, 1977). Depending on the precise location of the lesion and lesion/tumor size, the dysfunction may affect limb movements (Fontaine, Capelle, & Duffau, 2002; Peraud, Meschede, Eisner, Ilmberger, & Reulen, 2002; Russell & Kelly, 2003; Zentner, Hufnagel, Pechstein, Wolf, & Schramm, 1996), or it may be restricted to speech (Krainik et al., 2003; Mendez, 2004a, 2004b; Pai, 1999a, 1999b). This constellation of symptoms is termed the “SMA syndrome” (Potgieser, de Jong, Wagemakers, Hoving, & Groen, 2014). However, unlike with lesion to the IFG, in most cases, the disorders are only transient, resolving within weeks to months (Bannur & Rajshekhar, 2000; Laplane et al., 1977; Potgieser et al., 2014), with days to recovery correlated with interhemispheric connectivity between the SMA and the primary motor cortex (Oda, Yamaguchi, Enomoto, Higuchi, & Morita, 2018; M. Vassal et al., 2017). In the case of speech recovery, recent evidence suggests that left hemisphere speech functions are relocated to the right hemisphere following resection (Chivukula, Pikul, Black, Pouratian, & Bookheimer, 2018). Together, these findings clearly indicate that regions connected via the FAT—namely left IFG and pre-SMA/SMA have an important role in oral language/speech, but they also suggest some potential mechanisms of compensation in the case of damage to the FAT.

Information about the role of the FAT has also come from studies of electrical stimulation of the FAT. Vassal and colleagues (2014) performed electrostimulation of the FAT in an awake right-handed participant during resection of a glioma impacting the left frontal lobe. Although no speech and language deficits were noted before the surgery, the researchers induced speech arrest upon stimulation of the FAT, with normalization of speech when stimulation was stopped. Fujii and colleagues (2015) conducted a similar study in five right handed patients with left frontal lobe tumors. The target of stimulation was verified to be the FAT by pre-operative DW-MRI tractography. Speech arrest upon stimulation was observed in four out of five cases, with speech initiation delay also reported in one case. Finally, in a much larger study, Kinoshita and colleagues (2015) investigated 19 patients with frontal lobe tumors (14 left and 5 right). In sixteen of these participants, intraoperative electrostimulation of the FAT resulted in speech inhibition (arrest). Postoperative disturbances in speech, however, were limited to cases in which the left FAT was impacted, and no cases of speech disturbance were reported for right FAT lesion.

The left FAT is also associated with persistent developmental stuttering (also known as stammering; Kemerdere et al., 2016). Stuttering is characterized by disordered verbal fluency that appears in childhood and continues into adulthood. There is a lack of consensus about whether stuttering is primarily a disorder of language (Bernstein Ratner, 1997) or of motor coordination (Ludlow & Loucks, 2003; Max, Guenther, Gracco, Ghosh, & Wallace, 2004; Namasivayam & van Lieshout, 2011). There is also debate about the key brain regions associated with stuttering (Etchell, Civier, Ballard, & Sowman, 2018)—indeed fluent speech requires the recruitment of large-scale bilateral neural regions (Crinion, 2018). However, meta-analyses suggest several “neural signatures” of stuttering. These include abnormalities of the SMA, cerebellum, auditory cortex, basal ganglia, and right frontal operculum/insula regions (Brown, Ingham, Ingham, Laird, & Fox, 2005; Budde, Barron, & Fox, 2014; Watkins, Smith, Davis, & Howell, 2008). But stuttering research also routinely implicates the left and right IFG/PMv (Neef et al., 2016). The IFG/PMv area has been shown to be functionally and structurally anomalous (Watkins et al., 2008) and underactivated (Budde et al., 2014; Neef et al., 2016) in people who stutter. Recently, Chesters and colleagues (Chesters, Mottonen, & Watkins, 2018) targeted the left IFG/PMv using transcranial direct current stimulation (tDCS) and found improved speech fluency in people who stutter. The implication of the IFG in stuttering suggests a potential role for the FAT in this disorder.

There is still, though, a lack of consensus on the fiber pathway systems associated with stuttering (Ingham, Ingham, Euler, & Neumann, 2017; Kronfeld-Duenias, Amir, Ezrati-Vinacour, Civier, & Ben-Shachar, 2016a, 2017; Neef, Anwander, & Friederici, 2017). Structural differences are reported in the white matter underneath the IFG, angular gyrus, premotor cortex, and middle frontal gyrus (Chang, Zhu, Choo, & Angstadt, 2015; Connally, Ward, Howell, & Watkins, 2014; Watkins et al., 2008), and in the anterior segment of the right arcuate fasciculus (Kronfeld-Duenias et al., 2016a). Three recent studies, however, have specifically focused on the involvement of the FAT in persistent developmental stuttering. In the first study, Kronfeld-Duenias and colleagues (Kronfeld-Duenias, Amir, Ezrati-Vinacour, Civier, & Ben-Shachar, 2016b) examined 34 adults (15 of whom were stutters with a history of stuttering since childhood). Mean diffusivity of the left FAT differed between adults who stutter and controls, and predicted individual differences in speech rate in the individuals who stutter, which was interpreted as supporting evidence that the FAT is part of a “motor stream” for speech, as proposed by Dick and colleagues (2014). In the second study of eight patients undergoing surgery for glioma, Kemerdere and colleagues (Kemerdere et al., 2016) showed that transient stuttering can be induced via direct electrical stimulation of the left FAT during awake surgery. Furthermore, in cases where the FAT was spared from resection, patients experienced no post-operative stuttering. Finally, Neef and colleagues (Neef et al., 2018; Neef et al., 2016) investigated 31 adults with stuttering and 34 controls. They found that more severe stuttering was linked to stronger connectivity of the right FAT, which they interpreted as reflecting an overly active global motor inhibition commonly associated with the right inferior frontal gyrus. In sum, although additional research is needed, current evidence across modalities suggests that the FAT may be an important pathway for speech fluency in people who stutter.

The FAT has also been associated with verbal fluency performance more generally. Verbal fluency tasks typically require a participant to produce words beginning with a particular letter (e.g., “f”), or which come from a particular category (e.g., “animals”). The former are typically referred to as phonological fluency tasks, and the latter as semantic or category fluency tasks. Both tasks recruit the left IFG (Costafreda et al., 2006; Smirni et al., 2017) and the pre-SMA/SMA (Abrahams et al., 2003; Alario et al., 2006; Crosson et al., 2001; Persson et al., 2004; Tremblay & Gracco, 2006; Ziegler, Kilian, & Deger, 1997). Connectivity between the left IFG and pre-SMA/SMA could thus be expected to support the function of establishing a preferred motor response in the linguistic domain, and there is evidence that this is the case. For example, Kinoshita and colleagues (2015) reported an association between the distance from the FAT of the tumor resection and scores on post-operative semantic and phonemic fluency. In a study of patients with primary progressive aphasia (PPA), Catani and colleagues (2013) found that microstructural properties of the FAT, as measured by DW-MRI, were correlated with mean length of utterance and word per minute scores. In another study of PPA patients, FA of the left FAT was associated with speech fluency impairments (specifically number of speech distortions, speech rate, and syntactic production; Mandelli et al., 2014). Speech fluency was also related to FAT fractional anisotropy (FA), but only in the right hemisphere, in a sample of 10 minimally verbal children with autism (Chenausky, Kernbach, Norton, & Schlaug, 2017). Finally, Li and colleagues (2017) used DW-MRI and lesion-symptom mapping to study 51 right-handed stroke patients to determine which fiber pathways are associated with semantic and phonemic fluency. Semantic and phonemic fluency were negatively associated with lesion of the left FAT, and positively associated with FA of the left FAT.

In a controlled case study that is particularly illustrative of potential FAT function with respect to fluent speech, Chernoff and colleagues (Chernoff et al., 2018) examined two patients who underwent pre- and post-operative imaging and testing of speech and language function. The first patient underwent surgery for a left frontal glioma, which caused selective reduction in DW-MRI and fMRI metrics of connectivity of the left FAT. This patient experienced selective impairment in speech production with no impairment in lexical access. The second patient, who underwent left hippocampal resection, had no difficulty with spontaneous speech. Neither patient had difficulty postoperatively with word reading, nonverbal semantic processing, praxis, or motor function. Notably, the first patient’s difficulty with fluent speech was restricted to voluntary speech fluency of more complex sequences—the patient had no difficulty with sentence repetition or with picture naming. The patient also had no general or speech motor deficit.

The left IFG is also associated with a number of other more componential linguistic functions, including controlled lexical and phonological selection/retrieval (Badre et al., 2005; Devlin et al., 2003; Gough et al., 2005; Katzev et al., 2013), syntactic processing (Friederici, Ruschemeyer, Hahne, & Fiebach, 2003; Love, Swinney, Walenski, & Zurif, 2008), and production of speech and language more broadly (Guenther, 2016). It is not surprising, therefore, to expect that the FAT might be associated with these linguistic processes, and there is some evidence for this. For example, in young children the length of the left FAT predicts receptive language abilities (Broce, Bernal, Altman, Tremblay, & Dick, 2015). Some compelling evidence is presented by Sierpowska et al. (2015). In this case study of a patient undergoing resection for left frontal tumor, these authors showed that intraoperative stimulation of the left FAT elicited word retrieval deficits in a noun-verb morphological derivation task. When asked to generate a verb associated with a noun (e.g., book), the patient extended a morphological rule to invent a new word (e.g., booked) instead of producing an appropriate existing word (e.g., read). Notably, the patient did not display more extended verbal fluency deficits—in this case the deficit was specific to morphological derivation. Catani and colleagues (Catani et al., 2013) found a similar association with syntactic function and the FAT. Abnormality of the tract was most associated with the non-fluent/agrammatic subtype of PPA. Furthermore, the association between FA of the FAT and performance on an anagram test, a measure of grammatical processing, was *r* = 0.49, *p* = .03 (although this did not survive correction for multiple comparisons). FA of the left FAT was similarly associated with syntactic production scores in another sample of patients with PPA (*r* = 0.76, *p* = .02; Mandelli et al., 2014). These studies provide initial suggestive evidence for a functional association between the left FAT and syntactic processing.

### Summary

Much of the work on the associated functions of the FAT has been conducted with the aim to define its relation to speech and language functions. The extant data suggest that the tract is strongly associated with speech initiation, verbal fluency, and stuttering. Some initial associations have been made between the tract’s microstructure and higher-level language functions. Additional data will serve to further identify the specific linguistic functions of the pathway.

## 3. Functional Associations of the Right Frontal Aslant Tract in Executive Function/ Inhibitory Control

Although speech and language functions are distributed across several regions on both hemispheres, some aspects of speech and language are left lateralized in most right-handed individuals (Knecht et al., 2000), and the function of the left IFG has been a focus of inquiry since the time of Broca. The functional association of the right IFG has only more recently become a subject of debate. Earlier reports focused on the role of the right IFG in executive function, specifically inhibitory control/stopping behaviors—i.e., countermanding an initiated response tendency via top-down executive control, recruited during Go/NoGo and Stop-Signal experimental paradigms. In these tasks, a prepotent response is initiated (a Go process) that must be over-ridden when a stop-signal occurs (the Stop process; (Aron, 2007; Aron, Fletcher, Bullmore, Sahakian, & Robbins, 2003; Aron, Monsell, Sahakian, & Robbins, 2004). The initial evidence for the role of right IFG in inhibitory control came from neuroimaging studies using these tasks (Bunge, Dudukovic, Thomason, Vaidya, & Gabrieli, 2002; Garavan, Ross, & Stein, 1999; Konishi et al., 1999; Konishi, Nakajima, Uchida, Sekihara, & Miyashita, 1998; Menon, Adleman, White, Glover, & Reiss, 2001) and studies in patients with right inferior frontal cortex lesions (Aron et al., 2003; Aron et al., 2004). In these latter lesion studies, injury to the right IFG (specifically the IFGOp) was associated with inhibitory control and impaired inhibition of irrelevant task sets.

More recent research has focused on a more extended network implementing inhibitory control, including the dorsolateral prefrontal cortex, pre-SMA, SMA, dorsal anterior cingulate, supplementary eye field, frontal eye field, subthalamic nucleus, globus pallidus, and thalamus (Aron, 2007; Aron, Herz, Brown, Forstmann, & Zaghloul, 2016; Aron & Poldrack, 2006; Chambers, Garavan, & Bellgrove, 2009; Fife et al., 2017; Garavan et al., 1999; Jahanshahi, Obeso, Rothwell, & Obeso, 2015; Levy & Wagner, 2011; Wiecki & Frank, 2013). The outcome of the network interactions of these regions is proposed to be the suppression of cortical output for behaviors that conflict with a goal or target behavior (Wessel & Aron, 2017). The FAT, linking the inferior frontal and pre-SMA nodes, is an understudied connection in this network, but the evidence to which we will now turn suggests it is an important component (Vilasboas, Herbet, & Duffau, 2017).

In several models of inhibitory control (Aron & Poldrack, 2006; Wiecki & Frank, 2013), the right IFG directly activates neurons of the subthalamic nucleus through a direct pathway, which plays an explicit role in stopping motor behavior (Cai & Leung, 2009; Favre, Ballanger, Thobois, Broussolle, & Boulinguez, 2013; Frank, 2006; Jahanshahi, 2013; Obeso et al., 2014; van Wouwe et al., 2017). Thus, the early suggestion has been that suppression occurs through a direct interaction with right IFG and subthalamic nucleus. However, there is also suggestion that this connection proceeds through the pre-SMA (Aron et al., 2016).

In fact, this is consistent with the role of right pre-SMA and SMA in motor control more broadly, and in stopping behaviors in particular (Nachev, Kennard, & Husain, 2008). For example, fMRI studies show the right pre-SMA is more active when participants successfully stop a behavior compared to when they don’t (Aron et al., 2007; Aron & Poldrack, 2006; Boehler, Appelbaum, Krebs, Hopf, & Woldorff, 2010), and some have argued that this pre-SMA activation is a signature of successful inhibition (Sharp et al., 2010). Indeed, direct stimulation of both the right IFG and the right SMA/pre-SMA stops the production of ongoing movements (Luders et al., 1988; Mikuni et al., 2006), and right pre-SMA specifically activates in situations in which a participant must choose to perform a new response in favor of an established response (Garavan, Ross, Kaufman, & Stein, 2003). Right SMA/pre-SMA lesion impairs production of complex sequenced movements for both the contralesional and ipsilesional side of the body (J. P. Dick, Benecke, Rothwell, Day, & Marsden, 1986) and the resolution of conflict between competing action plans (Nachev, Wydell, O’Neill, Husain, & Kennard, 2007). Temporary lesion via transcranial magnetic stimulation (TMS) of the right pre-SMA also impairs stopping, resulting in longer response times in the stop signal paradigm (Cai, George, Verbruggen, Chambers, & Aron, 2012). Finally, a rare study implementing single-unit recording of an awake human shows pre-SMA neurons appear to play a role in the selection and preparation of movements (Amador & Fried, 2004). Thus, right pre-SMA and SMA seem to be important for the implementation of inhibitory control.

In such a role, the pre-SMA and SMA may determine response threshold directly through interactions with M1 (Chen, Scangos, & Stuphorn, 2010), or by influencing inhibitory and excitatory outputs of the basal ganglia back to cortex in a task-dependent manner (Aron et al., 2016; Bogacz, Wagenmakers, Forstmann, & Nieuwenhuis, 2010; Frank, 2006; van Veen, Krug, & Carter, 2008; Wiecki & Frank, 2013). However, this likely occurs within the context of interactions with right IFG—indeed, both regions are consistently active when preparing to stop and during stopping (Chikazoe et al., 2009; Zandbelt & Vink, 2010). The nature of these interactions has been studied in a patient with ECoG electrodes implanted over both the right pre-SMA and right IFG, from which recordings were made during a stop-signal task (Swann et al., 2012). In that study, it was shown that coherence between right pre-SMA and right IFG increased for stop-signal trials, suggesting that these regions make up a physiologically connected circuit engaged during tasks requiring stopping/inhibitory control. Swann et al. also identified, using DW-MRI, that these regions are structurally connected via the FAT (although at the time they did not explicitly label the pathway as the FAT).

### Summary

The emerging evidence suggests that interactions between right IFG and pre-SMA/SMA could be important for inhibitory control. This may be because right IFG is a locus of inhibitory control directly and communicates with pre-SMA/SMA and with subcortical basal ganglia structures (Aron, Robbins, & Poldrack, 2014), or it may be because the right IFG functions in controlled context monitoring, and activates in response to detection of salient targets, thereby influencing activity in pre-SMA/SMA (Chatham et al., 2012; Erika-Florence, Leech, & Hampshire, 2014; Hampshire, 2015; Hampshire, Chamberlain, Monti, Duncan, & Owen, 2010). From either perspective, the right FAT is a potential fiber pathway supporting inhibitory control.

## 4. Proposed Function of the Frontal Aslant Tract

As Schwan and colleagues (2012) point out, the limited physiologic data on this particular connection in humans makes it difficult to specifically determine its function at a more mechanistic level. Yet, several groups have hinted at proposed functions of the FAT, especially for its general involvement in speech. For example, Catani and Bambini (2014) proposed a broad role for the FAT in providing “a basis for intentional communicative acts”, with a potential role in social cognition. However, this is a rather coarse description of the function of the tract—the FAT certainly may be involved in intentional communication and social cognition, but these are rather broad functions. Here, we attempt to provide a systematic proposal for the function of the FAT based on the evidence that is available. Though admittedly speculative given the limited data, we believe this is important to help guide the much-needed research on this fiber pathway.

Stated simply, our proposal is that the FAT is a key component of a cortico-basal ganglia-thalamic-cerebellar circuit involved in action control. This circuit, with the FAT highlighted, is illustrated in Figure 2. More specifically, based on the evidence reviewed above, the FAT is best described as a pathway involved in the planning, timing, and coordination of sequential motor movements, and in the resolution of competition among possible voluntary sequential motor movements. Compelling empirical evidence for the *voluntary* function of the FAT is the association between resection of the FAT and incidence of the transient Foix-Chavany-Marie syndrome, which describes the loss of the voluntary control of facial, lingual, pharyngeal, and masticatory musculature in the presence of preserved reflexive and automatic functions of the same muscles (Brandao, Ferreria, & Leal Loureiro, 2013; Martino, de Lucas, Ibanez-Plagaro, Valle-Folgueral, & higher-level function resolving conflict among competing motor programs, thus establishing a directed movement.

**Figure 2:**
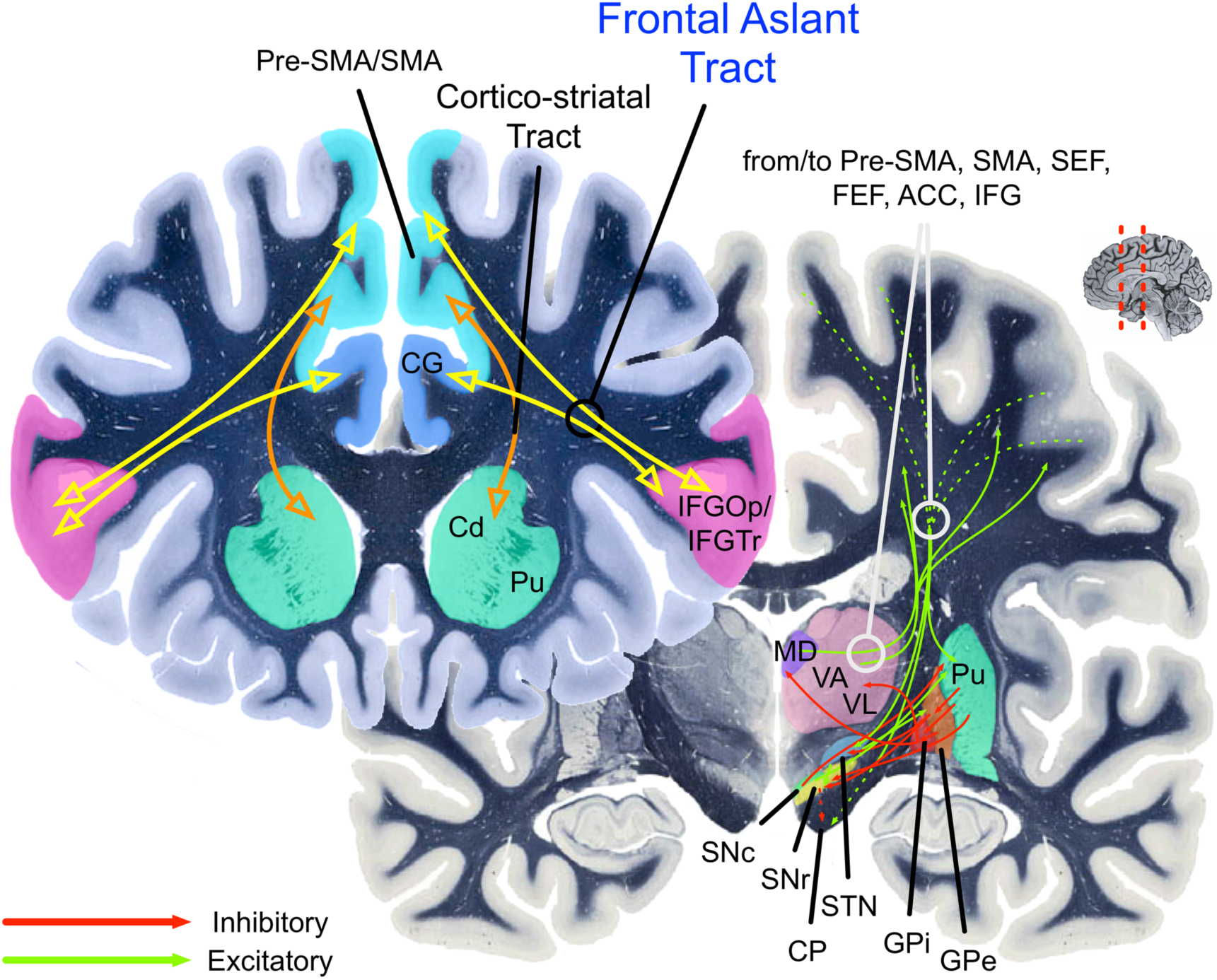
The cortico-basal ganglia circuits involved in speech production (left lateralized) and inhibitory control (right lateralized). The yellow path indicates the Frontal Aslant Tract (FAT). The orange path indicates the cortico-striatal tracts. Pre-SMA = pre-supplementary motor area; SMA = supplementary motor area; STN = subthalamic nucleus; thalamus VA/VL = ventral anterior and ventral lateral thalamic nuclei; GPe and GPi = globus pallidus external and internal; SNc = substantia nigra, *pars compacta*; SNr = substantia nigra, *pars reticulata*. CP = Cerebral peduncle. Pu = Putamen. Cd = Caudate. MD = Medialdorsal nucleus of thalamus. CG = Cingulate gyrus. ACC = Anterior cingulate cortex. SC = superior colliculus. Green arrows indicate excitatory connections. Red arrows indicate inhibitory connections. Hashed green arrows indicate originating fibers from multiple cortical areas arriving together in the internal capsule as they pass to their targets.

The function of the FAT and its involvement in resolution of competition among competing motor plans is proposed to be the same across the two hemispheres. However, here we propose some domain specialization across hemispheres. On the left hemisphere, this circuit is specialized for speech actions, although it may also participate in manual movements (Budisavljevic et al., 2017). On the right hemisphere, this circuit is specialized for general action control mechanisms, especially in the visuo-spatial domain. In both cases, the FAT plays a role in selecting among competing representations for actions that require the same motor resources (mainly the articulatory apparatus on the left hemisphere, and the oculomotor and manual/limb action systems on the right hemisphere).

The first piece of evidence in favor of this proposal is that the cortico-basal ganglia-thalamic-cerebellar circuits for speech and for oculomotor and manual/limb actions involve essentially homologous regions across the hemispheres (Figure 3 shows the cortical activations that are relevant for the FAT). This is in keeping with the established phenomenon that cortico-subcortical loops share a similar computational and broadly-defined architecture, but differ in terms of their specific connectivity with the originating and terminating cortical areas, and subregions of the striatum, cerebellum, and thalamus (Middleton & Strick, 2000).

**Figure 3:**
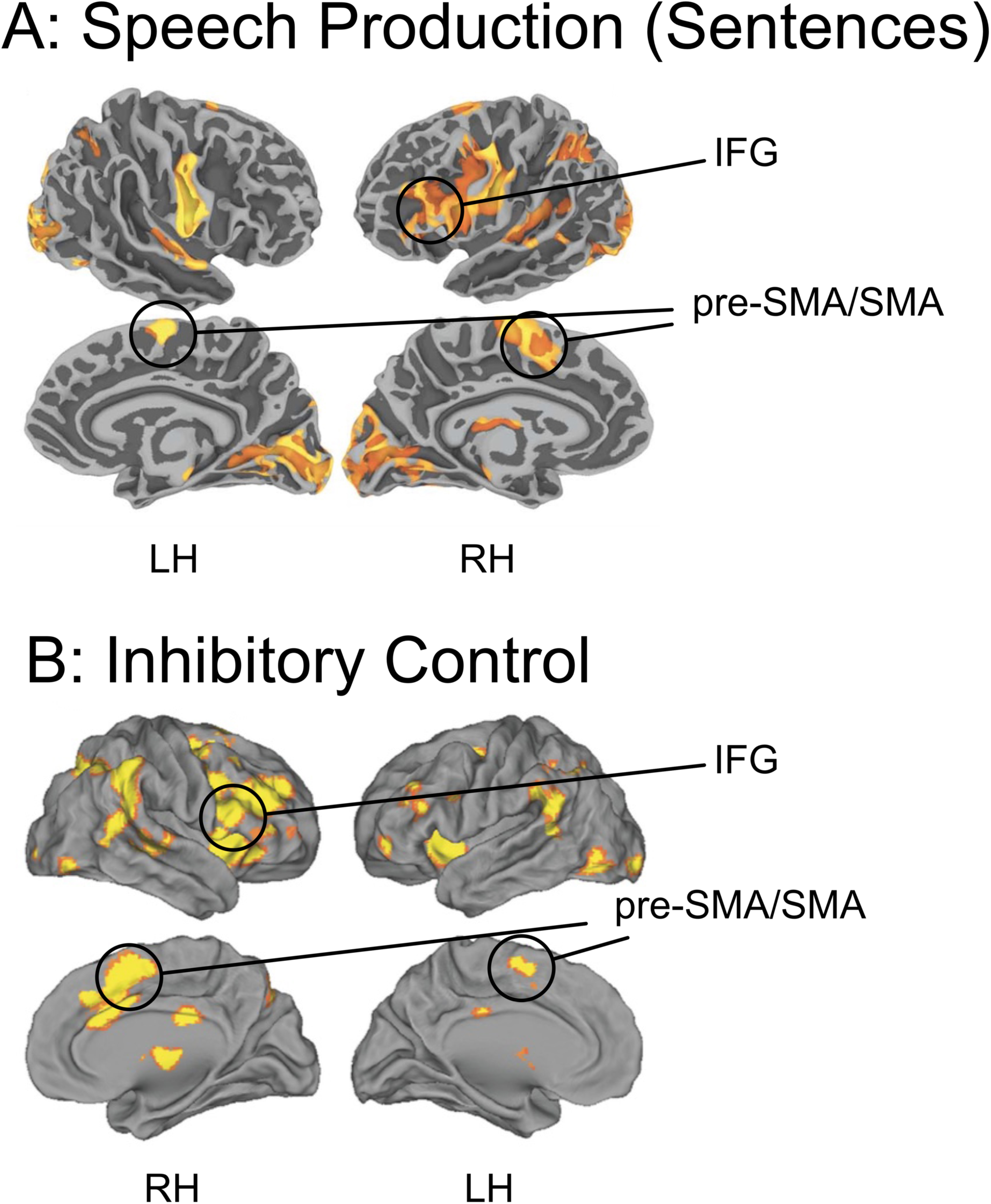
Top: Blood Oxygenation Level Dependent (BOLD) activation (relative to resting baseline) for sentence production in response to visual prompts (i.e., generating sentence from object pictures; IFG = inferior frontal gyrus; pre-SMA/SMA = pre-supplementary motor area/supplementaty motor area; from Tremblay, P., & Small, S. L. (2011). Motor response selection in overt sentence production: a functional MRI study. *Front Psychol, 2*, 253. Bottom: Results from a meta-analysis of inhibitory control from Cai, W., Ryali, S., Chen, T., Li, C. S., & Menon, V. (2014). Dissociable roles of right inferior frontal cortex and anterior insula in inhibitory control: evidence from intrinsic and task-related functional parcellation, connectivity, and response profile analyses across multiple datasets. *J Neurosci, 34*(44), 14652-14667. Studies included in the meta-analysis were conducted on healthy adults, using the stop-signal or go/no-go task with manual responses.

Side-by-side comparisons of computational models of speech and inhibitory control, which have to-date been developed largely independently, also illustrate this. Figure 4 (left) shows an influential computational model of speech, the Directions Into Velocities of Articulators (DIVA) model, and its extension to account for multisyllabic planning, the Gradient Order DIVA (GODIVA) model, proposed by Guenther (Bohland, Bullock, & Guenther, 2010; Guenther, 1992, 1994, 1995, 2016; Guenther, Ghosh, & Tourville, 2006; Guenther, Hampson, & Johnson, 1998). In this model, activation of a “cognitive context” of abstract phonemic and syllable frames, represented in the posterior inferior frontal sulcus and in pre-SMA, facilitates interactions between the basal ganglia and the SMA to initiate a specific speech motor program. Essentially, in this model, the basal ganglia establish a “winner-take-all” competition between competing speech motor programs, with an initiation signal sent to SMA when the cognitive, motor, and sensorimotor patterns match the context of a particular specific motor program. The extended GODIVA model provides for an inhibitory connection between the left posterior IFS and the left pre-SMA, which are activated in parallel. This interaction supports “winning” potential phonemes (represented in IFS) and syllable frames (represented in pre-SMA). Although not explicitly stated, it can readily be hypothesized that that functional interactions between IFS and pre-SMA are structurally supported by the FAT.

**Figure 4:**
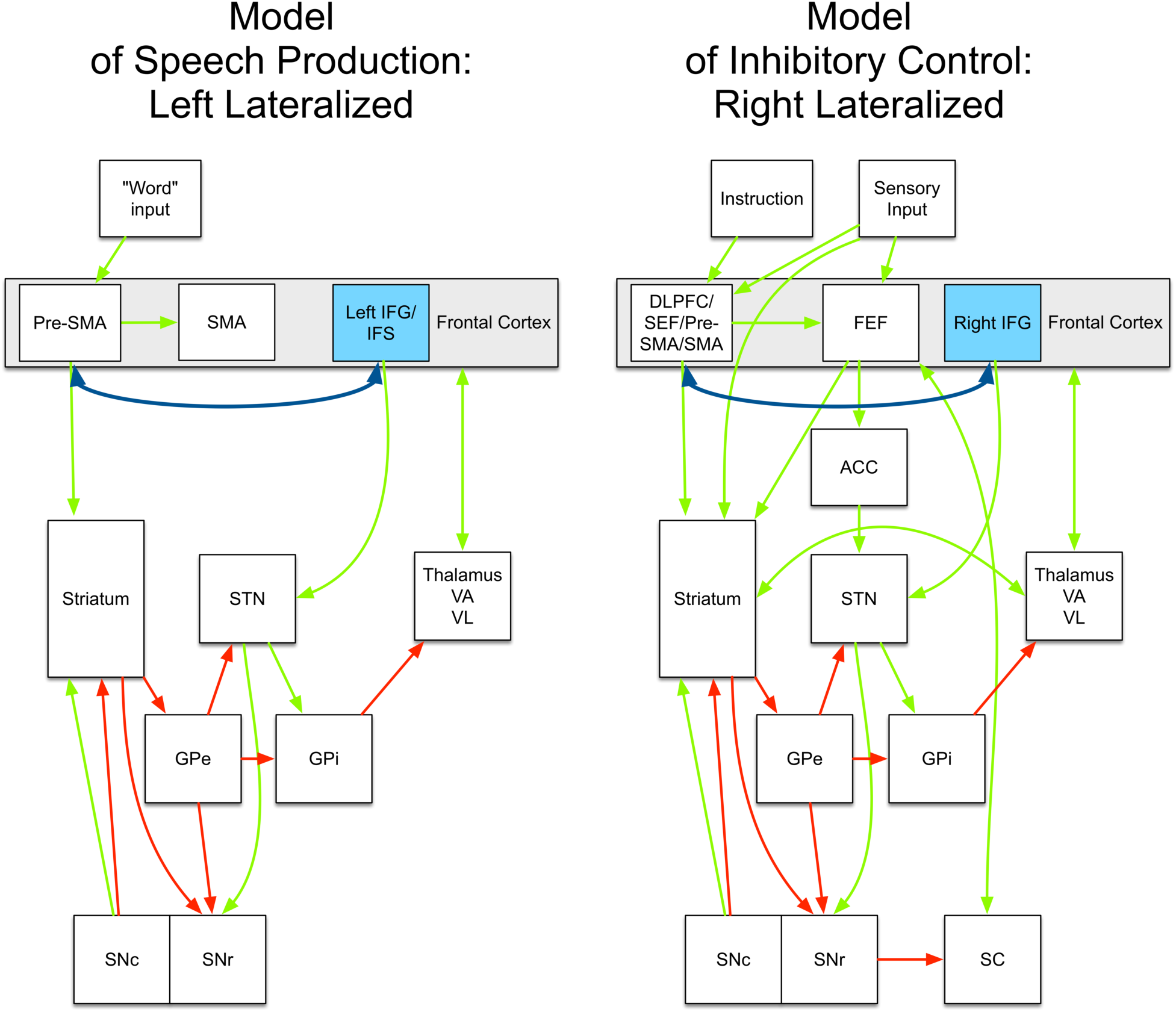
Left: An example connectivity model of speech production, from Guenther (Guenther, 2016). For simplicity, the model is incomplete (e.g., it does not include the cerebellum). The complete model is specified in Guenther (2016). Right: An example connectivity model of inhibitory control, based on Wiecki and Frank (Wiecki & Frank, 2013). The dark blue double-headed arrow represents the frontal aslant tract (FAT). Pre-SMA = pre-supplementary motor area; SMA = supplementary motor area; STN = subthalamic nucleus; thalamus VA/VL = ventral anterior and ventral lateral thalamic nuclei; GPe and GPi = globus pallidus external and internal; SNc = substantia nigra, *pars compacta*; SNr = substantia nigra, *pars reticulata*. ACC = Anterior cingulate cortex. SC = superior colliculus. Green arrows indicate excitatory connections. Red arrows indicate inhibitory connections.

While DIVA/GODIVA focuses on phonemic and syllable-level representations, the data reviewed above suggest that the FAT might also be involved in lexical-level retrieval and selection. This may involve the facilitation of interactions between more anterior IFG, proposed to be involved in semantic selection and retrieval (Badre et al., 2005; Devlin et al., 2003; A. S. Dick et al., 2014; Gough et al., 2005; Katzev et al., 2013), and the pre-SMA and SMA involved in establishing appropriate motor programs for speech. The left-lateralized FAT would presumably establish these action plans based on left-lateralized linguistic representations, codified in network-level interactions with left posterior temporal cortex.

Models of inhibitory control for manual actions propose a similar, but right lateralized, architecture. For example, the computational model proposed by Frank and Wiecki (Badre & Frank, 2012; Frank, 2006; Frank & Badre, 2012; Wiecki & Frank, 2013) establishes essentially the same basal ganglia loop comprising the direct and indirect pathways of known basal ganglia connectivity (Figure 4, right). In this model, specified for both manual responses and saccadic eye-movements (hence the inclusion of the superior colliculus), these basal ganglia connections implement selective gating of candidate motor actions (e.g., either a “Go” or “NoGo” action). The candidate actions, though, are determined by activity in frontal cortical regions. In the model specified by Wiecki and Frank (2013), rule-based representations are implemented in the right dorsal and lateral prefrontal cortex (DLPFC), supplementary eye-field (SEF), and pre-SMA. The pre-SMA is proposed to play a specific role in transforming the abstract rule representation into concrete candidate actions. The right IFG, however, applies a hyperdirect connection to the STN to facilitate a global stopping mechanism. No accommodation for connectivity between the right IFG and right pre-SMA is applied in this model. Consistent with this model, Aron et al. (2016) suggest that the right IFG and pre-SMA are part of dissociable circuits. The right IFG is proposed to be part of a fronto-STN pathway for stopping, while the pre-SMA is part of fronto-STN circuit for resolving conflict. However, the empirical data we reviewed above suggest that the direct connectivity between right IFG and pre-SMA *is* a potentially important component of the neural network implementing inhibitory control processes.

We suggest that the right IFG-pre-SMA connection supported by the FAT plays a similar role as it does in the left-lateralized network implementing speech. That is, broadly defined, interactions between these regions establish action plans for the output of sequential (non-speech) motor programs, and together decide on a “winning” action plan, implemented via downstream interactions in basal ganglia and motor cortex. Consistent with this idea are studies that show that diffusion FA of the right FAT is related to deficits in constructional apraxia (Serra et al., 2017), and to more efficient visuomotor processing during manual movements, resulting in smoother movement trajectories (Budisavljevic et al., 2017). This again points to the importance of this tract in sequential movement planning.

Aron and others (Aron et al., 2014; Swann et al., 2012) have proposed the possibility that it is the degree of synchrony between right IFG, pre-SMA, and basal ganglia, and not necessarily their local activity, that determines whether inhibition of a motor response occurs. The proposal that the pre-SMA plays a general task configuration role and directly mediates right IFG function in stopping is consistent with the structural connectivity of the FAT, and the physiologic data suggesting that the pre-SMA activates before right IFG during stop trials (Swann et al., 2012).

In summary, based on the available evidence, we suggest that the FAT implements resolution of competition among conflicting motor programs to implement voluntary sequential movement, with some level of hemispheric specialization.

## 5. Limitations and Suggested Areas of Future Research

The proposed model suggests a number of potential avenues for future research. We will focus on a few here. First, with respect to the left FAT, although we have argued for an important but perhaps not primary role for the FAT’s in sequential movement planning for speech, we have grounded this on a limited empirical base investigating the tract’s specific functions. Very limited research has examined different sub-components of the FAT and their associated functions. Thus, it may be the case that IFGTr and IFGOp connections with the pre-SMA play somewhat different functional roles for speech. We have also not established that additional connections (e.g., with anterior insula) are functionally important for speech. More focused study of these subcomponents is necessary. Also necessary is the study of post lesion reorganization, to help understand why speech disorders in patients with SMA syndrome are only transient. It would be interesting to determine whether the similar structural architecture across the two hemispheres facilitates reorganization of function to the contralesional network.

Second, our model does not firmly establish the connection with language functions that have no explicit motor component, such as syntax. We have neglected to speculate broadly on this, as we await further empirical evidence. However, it is possible that the link between speech and syntax is their inherently sequential nature. More research on this specific link is also needed.

Third, with respect to the right FAT, we have focused on only two broadly defined executive functions, inhibitory control and controlled context monitoring. Executive function is itself a non-unitary, broadly defined construct, but little research has investigated the link between executive function and the FAT that we may be simply scratching the surface. Thus, it is possible that the FAT plays a role in planning defined more broadly, or in other broadly defined executive functions. For example, a recent study suggests that left laterality of the FAT is associated with greater attention problems in children, although this association was fully mediated by executive function as measured by parent and teacher ratings (Garic, Broce, Graziano, Mattfeld, & Dick, in press). In another recent study, Varriano and colleagues (2018) showed that more extensive connectivity with anterior superior frontal gyrus via the right FAT is associated with working memory performance in adults. More work in this area is needed to understand which aspects of executive function are associated with the microstructural properties of the FAT. In particular, it will be important to examine connectivity with the anterior cingulate, which has been associated with cognitive control (Lovstad et al., 2012).

Finally, the model we propose suggests that the FAT may be a target of clinical significance. The FAT could be targeted as a structure expected to show change in response to clinical intervention for disorders of speech and language (e.g., stuttering, aphasia), or for disorders associated with inhibitory control deficits (e.g., attention deficit hyperactivity disorder; ADHD). Presurgical mapping of the FAT, in cases of surgical resection, may also become increasingly important if the desire is to spare some of the functions we have identified here.

## 6. Conclusion

The available data suggest the existence of direct pre-SMA and IFG connections. We propose that this connection is a key pathway for two important functional circuits—speech and executive function/inhibitory control—that are typically examined separately but that rely on overlapping mechanisms. What this review shows is that cross-pollination of the models of these circuits may be beneficial for understanding each of them separately. In addition to understanding the basic circuitry of these seemingly-disparate functions, the current models may also inform neurosurgical interventions and, in turn, may stimulate the application of new pre-surgical mapping techniques. The models may also establish targets expected to respond to clinical intervention for disorders of speech and language, or of executive function. Because the pathway is only recently defined, there is a rich empirical landscape available to help us answer some of the critical questions about its associated functions.

## Acknowledgements

This work was supported by grants from the National Institutes of Mental Health (Grant R01MH112588 and R56MH108616 to P.G. and A.S.D), and from the National Institute for Drug Abuse (U01DA041156; salary support to A.S.D.). P.T is supported by a career award from the from the “Fonds de la Recherche en Santé du Québec” (FRQS, #35016). Conflict of Interest: None.

